# Biological adaptation under fluctuating selection

**DOI:** 10.1101/457762

**Authors:** Ming Liu, Dustin R. Rubenstein, Wei-Chung Liu, Sheng-Feng Shen

**Affiliations:** Biodiversity Research Center, Academia Sinica, Taipei, 11529, Taiwan; Department of Ecology, Evolution and Environmental Biology, Columbia University, New York, NY 10027 USA; Center for Integrative Animal Behavior, Columbia University, New York, NY 10027 USA; Institute of Statistical Science, Academia Sinica, Taipei 11529, Taiwan

## Abstract

Bet-hedging—an evolutionary strategy that reduces fitness variance at the expense of lower mean fitness—is the primary explanation for most forms of biological adaptation to environmental unpredictability. However, most applications of bet-hedging theory to biological problems have largely made unrealistic demographic assumptions, such as non-overlapping generations and fixed population sizes. Consequently, the generality and applicability of bet-hedging theory to real world phenomena remains unclear. Here we use continuous-time, stochastic Lotka-Volterra models to relax overly restrictive demographic assumptions and explore a suite of biological adaptations to fluctuating environments. We discover a novel “rising-tide strategy” that—unlike the bet-hedging strategy—generates both a higher mean and variance in fitness. The positive fitness effects of the rising-tide strategy’s specialization to good years can overcome any negative effects of higher fitness variance in unpredictable environments. Moreover, we show not only that the rising-tide strategy will be selected for over a much broader range of environmental conditions than the bet-hedging strategy, but also under more realistic demographic circumstances. Ultimately, our model demonstrates that there are likely to be a wide range of ways that organisms respond to environmental unpredictability.

---

Temporal fluctuation of environmental conditions is a universal feature in nearly every ecosystem on earth^1,2^. In fluctuating environments, the intensity and direction of natural selection is likely to vary unpredictably over time^3-5^. Almost all known biological adaptations to environmental fluctuation—as diverse as seed production in annual plants^6^, phenotypic polymorphisms in bacteria^7-9^, and altruistic behavior in social animals^10^—are typically summarized by a single, simple mechanism: the evolutionary bet-hedging strategy^1,11,12^. There are two general types of bet-hedging: conservative bet-hedging describes a consistent but low risk strategy or phenotypic investment, whereas diversified bet-hedging depicts the case when organisms spread risk by investing in different strategies or phenotypes^11,13^. Both forms of bet-hedging result in the same fitness consequence: reduced variance in fitness at the expense of a lower mean fitness. In other words, under either bet-hedging scenario, natural selection can act optimally by minimizing fitness variance rather than by maximizing mean fitness.

Although the bet-hedging principle describes many forms of behavioral adaptation under fluctuating selection, it cannot explain all of the ways in which animals cope with environmental uncertainty. For example, studies exploring thermal niche evolution at different temporal scales of environmental fluctuation have demonstrated that although greater long-term environmental variation (e.g. seasonal variation in temperature) favors the evolution of thermal generalists, short-term variation (e.g. daily temperature variation) has an opposite effect by selecting for thermal specialists^14,15^. Essentially, these models hint at the theoretical possibility of a specialist strategy in which individuals have a higher mean fitness while also having higher variance in fitness. Such an adaptive strategy to environmental fluctuation—which differs fundamentally from a bet-hedging strategy—can also be derived using the approximation for geometric mean fitness, which increases with higher arithmetic mean fitness and lower fitness variance^16-18^. However, optimality models based on geometric mean fitness rely on the restrictive assumptions of non-overlapping generations and infinite population sizes, both of which do not apply to most eukaryotic species. Indeed, fluctuation in population size—which when environmentally-driven is analogous to a population going through a bottleneck in bad years and an expansion in good years^19^—can have substantial effects on the strength and direction of selection in populations of finite size^20^. The strength of natural selection, which can be represented by the opportunity for selection (defined as the variation of relative fitness^21,22^), is greater when populations decrease in size but lower when populations expand. However, the assumption of non-overlapping generations—which is typically used in most models of this sort (e.g. the grain-size model^23^)—creates a distinction between within-and among-generation selection, and only applies to a very limited number of real world organisms such as some microbes^10,24^. As a consequence, identifying general rules of biological adaptation to environmental fluctuation—particularly for species with overlapping generations and finite population sizes—remain elusive.

To achieve a more comprehensive understanding of biological adaptation to environmental fluctuation that applies to organisms without having to evoke restrictive demographic assumptions that are biologically unrealistic for most animal species, we use continuous-time, stochastic Lotka-Volterra models to examine the impact of differential selective forces with varying population sizes, different temporal scales of environmental fluctuation, and distinctive patterns of generational overlap. We use a competitive Lotka-Volterra model as our basic framework because this approach restricts population size through competition, rather than externally setting an absolute boundary on population size. In other words, population size is dynamically regulated by the fitness of each strategy, which is in turn affected by environmental conditions and population size itself. Moreover, Lotka-Volterra models can be used to explore the process of natural selection, as in Moran models that incorporate birth and death processes^25^, without the unrealistic assumption of a fixed population size^26^. Fundamentally, our model always allows for competition to reduce the number of surviving strategies, yet the direction of selection may not be the same at each moment because environmental conditions fluctuate (i.e. fluctuating selection).

We begin by employing the generalist-specialist trade-off concept^15,27^ to explore the selection dynamics of a generalist bet-hedging strategy that persists in both good and bad years and a specialist strategy that instead favors good years (hereafter referred to as the “rising-tide strategy”). Specifically, the specialist rising-tide strategy is defined as an evolutionary strategy that has relatively high fitness in good years so that its population size and relative frequency in the population increase, but relatively low fitness in bad years such that its population size and relative frequency decrease (Fig. 1). Thus, unlike the bet-hedging strategy, which can have higher mean fitness but lower variance in fitness, the rising-tide strategy can have both a higher mean fitness *and* a higher variance in fitness (Fig.1b). Importantly, just as the generalist and specialist strategies represent two ends of a continuum, the bet-hedging and rising-tide strategies should also be viewed as two ends of an adaptive continuum to fluctuating environments. In other words, the bet-hedging and rising-tide strategies are endpoints along a continuum of potential adaptations to environmental fluctuation that differ only in their relationships between fitness mean and variance.

**Fig. 1.**
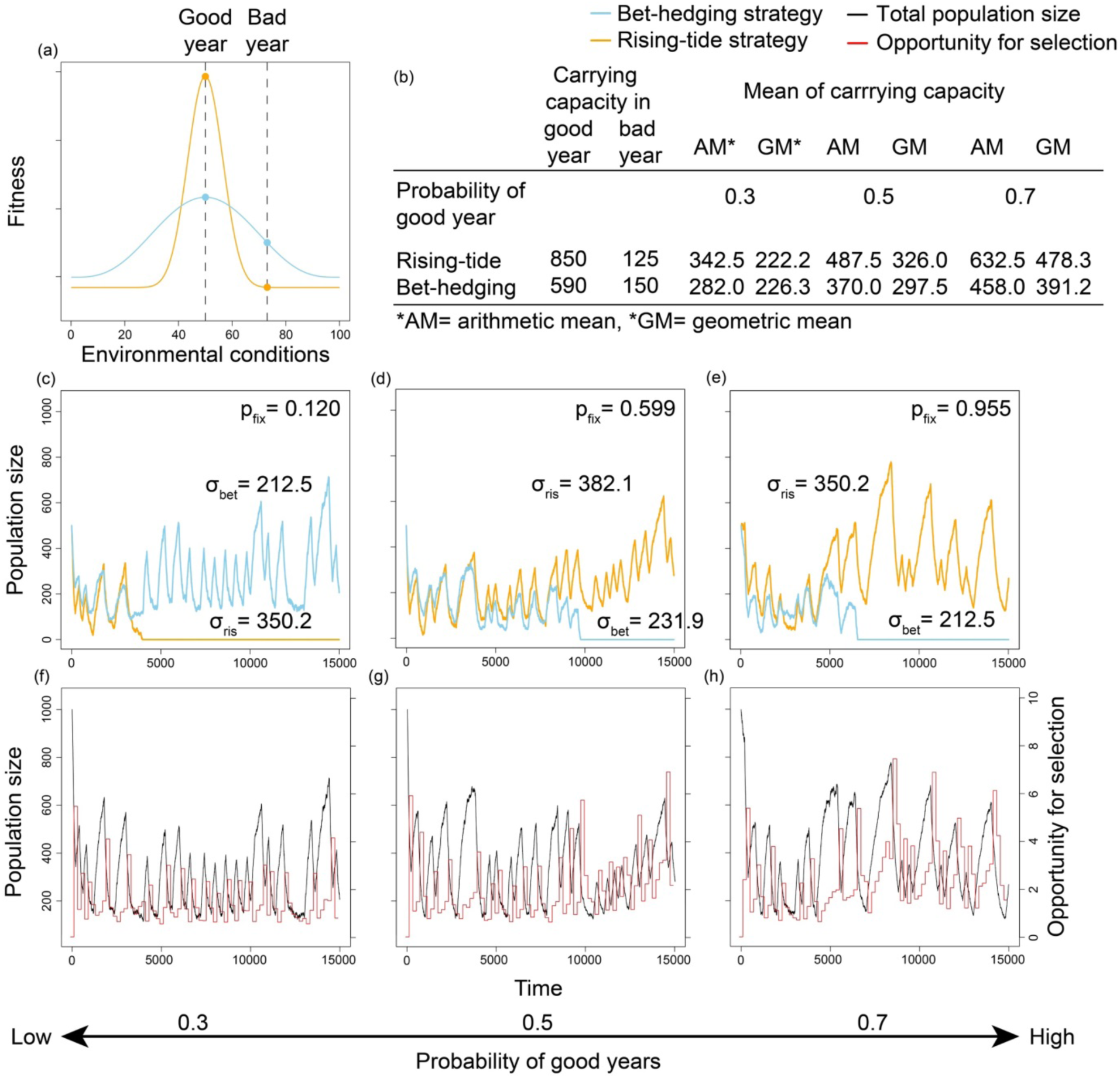
A comparison of the bet-hedging and rising-tide strategies under random environmental conditions and continuous population dynamics. **a,** Performance curves of the bet-hedging (blue line) and rising-tide strategies (orange line) under hypothetical environmental conditions, which can be thought to represent conditions like temperature or precipitation and is standardized from 0 to 100. These performances are modeled under discrete environmental conditions: (i) good years, where the environmental conditions are optimal for both strategies (i.e. environmental conditions = 50); and (ii) bad years, where the environmental conditions are non-optimal (i.e. environmental conditions = 75). **b,** These performances are expressed as the carrying capacities of a stochastic Lotka-Volterra model such that the rising-tide strategy has larger a carrying capacity in good years, and the bet-hedging strategy has a larger carrying capacity in bad years. Using these mean carrying capacities, we calculate the arithmetic (AM) and geometric means (GM) of fitness of a given probability of good years (i.e. environmental conditions = 50). **c-e,** We then explore the selection dynamics of the bet-hedging and rising-tide strategies (see Supplementary Table 1 for parameter values). The probability of fixation of the rising-tide strategy, as well as the fitness variations (σ) of the two strategies are shown in each panel. **f-h,** In the same time series as panel c-e, the intensity of selection in fluctuating environments can be described by the opportunity for selection (i.e. the variation of relative fitness, red lines), which is related to total population size (i.e. sum of the bet-hedging and rising-tide strategies; black lines). The value of opportunity for selection is updated once per year before the relative fitness of each individual is reset. Note that this model is a stochastic individual-based model and that a simplified deterministic model is presented in the supplementary information and Supplementary Fig. 1.

For simplicity, we begin by presenting the discrete version of the bet-hedging and rising-tide strategies under two types of environmental conditions: good and bad years (in a later section, we present the continuous versions of the generalist bet-hedging and specialist rising-tide strategies, as well as how they both respond to different temporal scales of environmental fluctuation). The selection dynamics of the rising-tide, *dN_R_*/*d_t_*, and bet-hedging, *dN_B_*/*d_t_*, strategies in good years are:

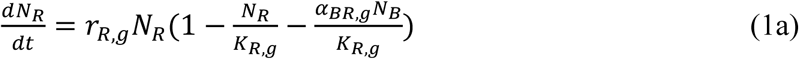

and

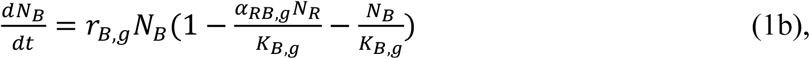

where *r* stands for the intrinsic growth rate (i.e. birth minus death rate), *N* represents the population size of each strategy, *α* describes the intensity of competition between two strategies, and *K*denotes the carrying capacity. The capital subscripts *R* and *B* represent the parameters for the *rising*-tide strategy and the *bet*-hedging strategy, respectively. Similarly, the lower-case subscripts *g* and *b* represent *good* and *bad* years, respectively. For example, *α_BR,g_* indicates the intensity of competition between the bet-hedging and rising-tide strategies in good years. Similarly, the selection dynamics of each strategy in a bad year are:

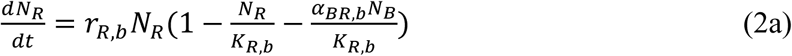

and

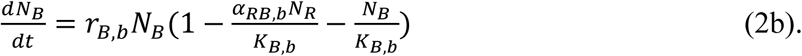

We begin by investigating the selection dynamics of the bet-hedging and rising-tide strategies using individual-based simulations, i.e. a stochastic continuous time model with random environmental settings, because this approach allows us to manipulate environmental stochasticity and track reproductive success at the individual level in order to calculate each strategy’s fixation probability and opportunity for selection (i.e. variance of relative fitness). We find that greater environmental variation can generate either the bet-hedging or the rising-tide strategy, but which end of the adaptive continuum is favored depends upon the frequency of good versus bad years (Fig. 1, see Supplementary Table 1 for parameter values). That is, the bet-hedging strategy has a higher fixation probability when bad years are more frequent than good years, whereas the rising-tide strategy is favored by selection when good years are more likely to occur than bad ones. When bad years are frequent, the risk aversion strategy of bet-hedging maximizes fitness by reducing variance rather than optimizing the mean, as has been shown previously^1,11-13^. However, when good years are more frequent than bad years, the positive impacts of a rising-tide strategy in the good years are able to sustain those individuals during the bad years (Fig. 1a-c). Importantly—and also consistent with previous theories—we find that the opportunity for selection rises when total population size declines (Fig. 1f-h) (results from the deterministic continuous time model with periodical environment setting are qualitatively similar; Supplementary Fig. 1). Therefore, the same magnitude of absolute fitness increase has a greater impact on the frequency of each phenotype in bad years than in good years. Thus, adaptation to bad years takes on a greater importance when bad years occur more frequently (and vice versa).

To further investigate how different temporal scales of environmental fluctuation—often referred to as the grain of the environmental variation; *sensu*^12^—influence the evolution of the bet-hedging and rising-tide strategies, we employ continuous versions of the generalist bet-hedging and specialist rising-tide strategies (Fig. 1a). Instead of assuming discrete environmental conditions (i.e. good versus bad years) as we did above, we now allow the environmental conditions *E* to vary continuously—just as temperature and rainfall do in nature—and influence the birth rate *b(E)*, death rate *d(E)*, and carrying capacity *K(E)* of each strategy (see Methods, eqns. 5 and 6). Thus, this is a stochastic continuous time model with random continuous environmental settings. The dynamics of the specialist rising-tide, *dN_R_*/*dt*, and generalist bet-hedging strategies, *dN_B_*/*dt*, in the stochastic Lotka-Volterra competitive model are:

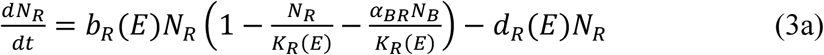

and

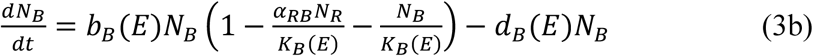

The capital subscripts *R* and *B* represent the parameters for *rising-tide* and *bet-hedging* strategy, respectively.

We find that long-term (e.g. annual) and short-term (e.g. daily) environmental fluctuations can have complex, but often counterintuitive, effects on adaptive evolutionary responses (Fig. 2, see Supplementary Table 2 for parameter values). When short-term environmental variation is relatively low, higher long-term variation selects for the bet-hedging strategy (Fig. 2c). However, the rising-tide strategy out-competes the bet-hedging strategy as short-term variation increases (Fig. 2d). When both short-and long-term environmental variation are relatively high, the bet-hedging strategy again becomes more dominant (Fig. 2e). Furthermore, when long-term environmental variation is relatively low, the rising-tide strategy is favored by selection under both low or high short-term environmental variation (Fig. 2i-k) because if long-term variation is relatively low, a rising-tide strategy, by definition, is more likely to specialize in a given mean environment (Fig. 2i). However, if long-term environmental variation is relatively high and if there is no, or only a small degree of, short-term environmental variation, by definition, a rising-tide specialist will not experience its optimal environmental conditions frequently (Fig. 2a and 2b). In other words, higher short-term variation can favor the evolution of specialization because a rising-tide specialist is likely to experience its optimal environmental conditions more frequently. This mechanism is similar to the one we described previously, namely that the specialist rising-tide strategy experiences more good than bad years and can therefore outcompete the generalist bethedging strategy, even when the variance in absolute fitness of the rising-tide strategy is higher than that of the bet-hedging strategy.

**Fig. 2.**
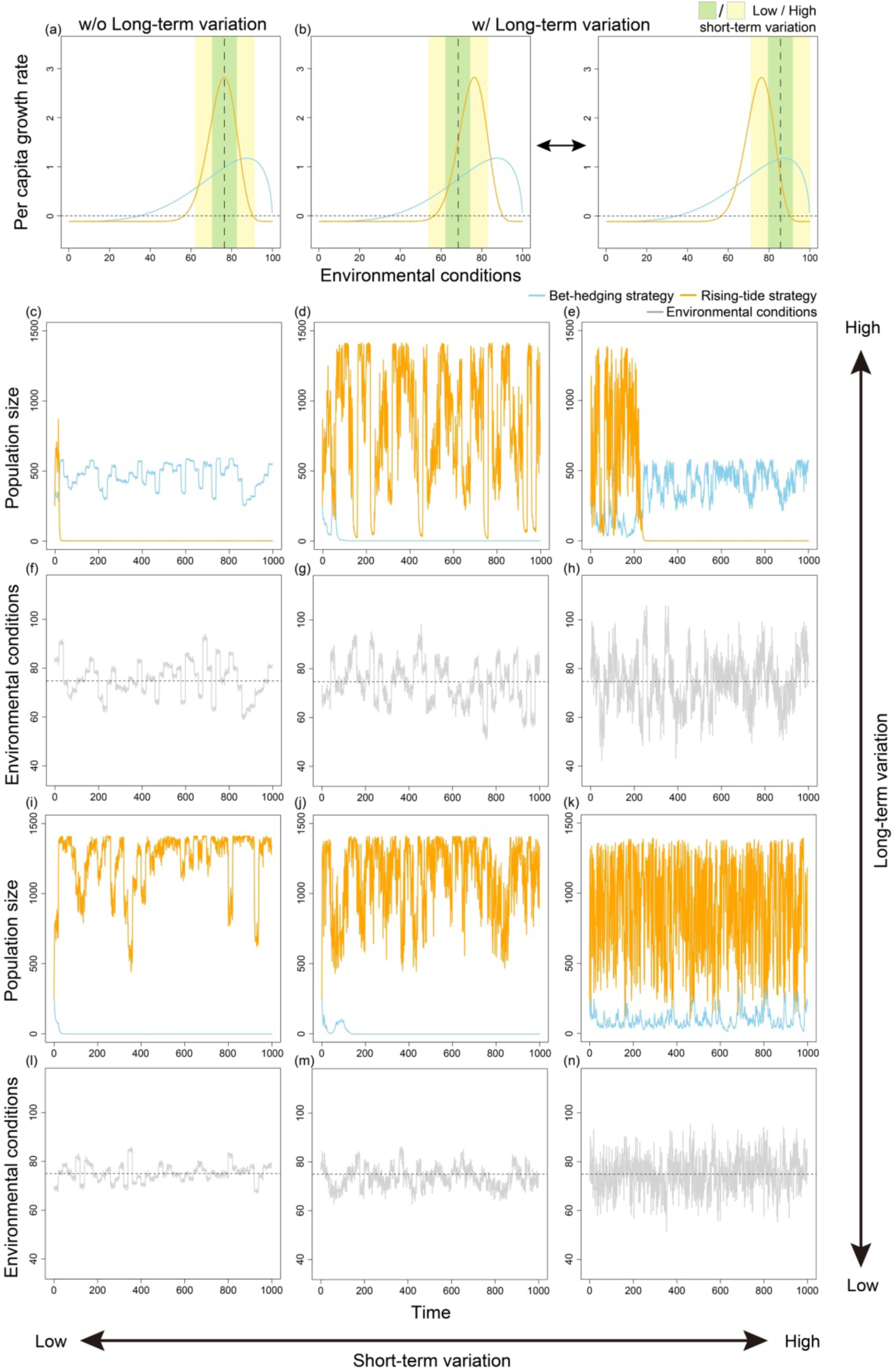
The selection dynamics of the bet-hedging and rising-tide strategies under random continuous environmental conditions with continuous population dynamics. **a,** Performance curves of bet-hedging (blue line) and rising-tide strategies (orange line) in hypothetical environmental conditions. Population dynamics parameters are functions of the environment, meaning average fitness (per capita growth rate) varies with changes in environmental conditions (see model description for more details). **b,** Potential effect of longterm environmental variation on fitness. Since long-term environmental variation may alter the mean of short-term environmental variation (thick dashed lines), low (green area) and high (yellow area) short-term variation can differentially alter the direction of selection. **c-e,** Population dynamics of the bet-hedging and rising-tide strategies under low, medium, and high short-term environmental variation, respectively, and under high levels of long-term environmental variation. **f-h,** Representations of the environmental conditions (i.e. time series) corresponding to panels **c-e. i-k,** Population dynamics of the bet-hedging and rising-tide strategies under low, medium, and high short-term environmental variation, respectively, and under low level of long-term variation. **l-n,** Representations of the environmental conditions (i.e. time series) corresponding to panels **i-k.** The parameter values for simulations are summarized in Supplementary Table 2, whereas the two environmental variables in this figure are *s_episode_* (17 and 100) and *s_phase_* (10, 100 and 1000).

To more directly determine how our framework performs relative to more traditional approaches, we compare our continuous time, overlapping generations model with one that utilizes discrete population dynamics to approximate the commonly used but less realistic non-overlapping generations models (e.g.^1,11,13,23^) (Fig. 3, see eqns. 7 and 8 in Methods for more details). In most of these models, the temporal scale of environmental fluctuation is often classified as either (i) coarse grain, which describes among-generation variation in environmental conditions, or (ii) fine grain, which describes within-generation variation in environmental conditions^12,23^. In our comparison, we find that patterns of generation overlap are crucial for the evolution of the generalist-specialist continuum and the adoption of a bet-hedging versus a rising-tide strategy (Fig. 4, see Supplementary Table 2 for parameter values). In this stochastic discrete time model with a random continuous environmental setting, when long-term variation is high—which is similar to coarse grain variation in discrete-population dynamics models—the bet-hedging strategy dominates the rising-tide strategy in most of the parameter space (Fig. 4a-h). Nevertheless, higher short-term variation—which is equivalent to fine grain variation in our model—still favors the rising-tide strategy when long-term variation is relatively low (Fig. 4i).

**Fig. 3.**
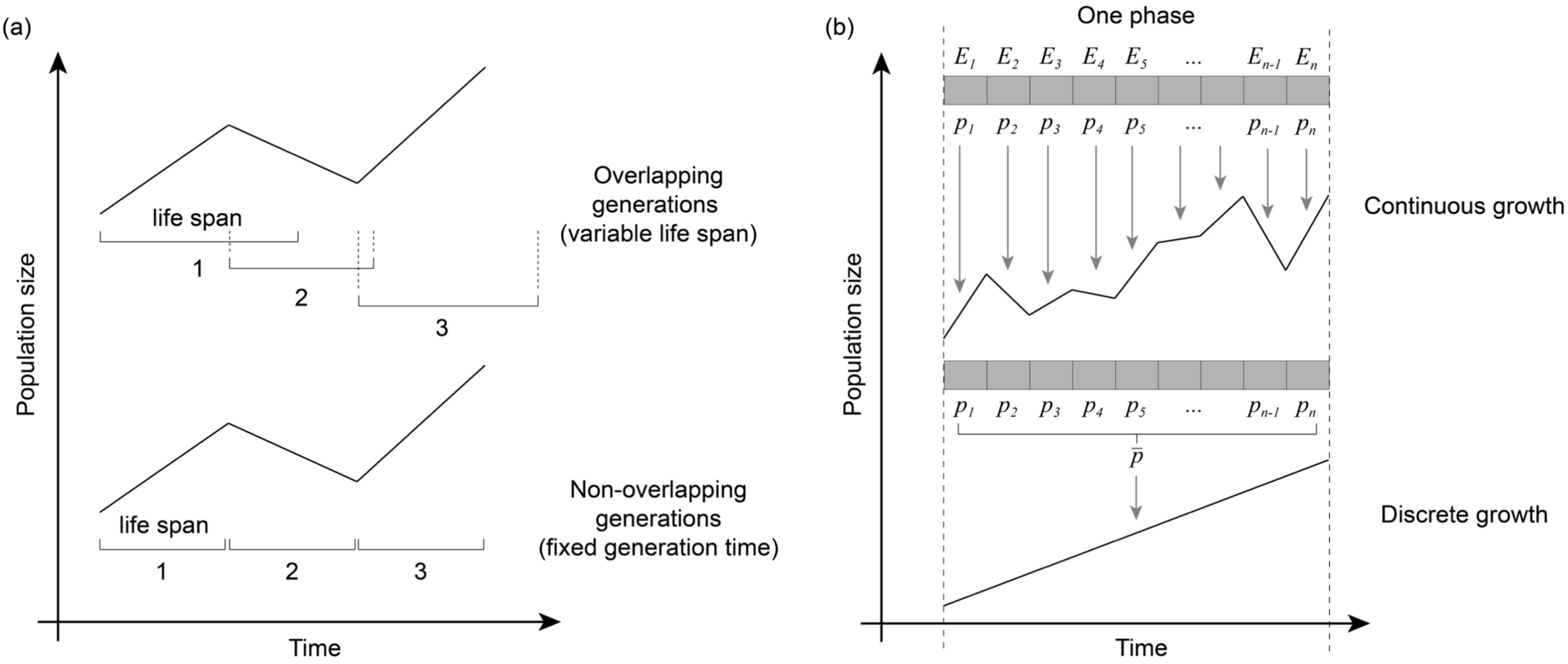
A schematic comparison of demographic growth settings. **a,** Although the overlapping and non-overlapping generation models may have similar population dynamics, the events within these dynamics are different. The main difference between the two models is that the overlapping generations model allows for lifespan (i.e. the numbers and brackets below population dynamics) changes with changing environmental conditions, whereas the non-overlapping generations model assumes fixed generation times. Moreover, the non-overlapping generations model assumes simultaneous death and birth events, which is hardly applicable to most, if not all, real organisms. **b**, Similar to the non-overlapping generations model, the discrete growth model takes the average (i.e. *p̅*) over many growth parameters (i.e. *p*) whenever a fixed time span has passed, whereas the continuous growth model applies each growth parameter instantaneously to the population dynamics.

**Fig. 4.**
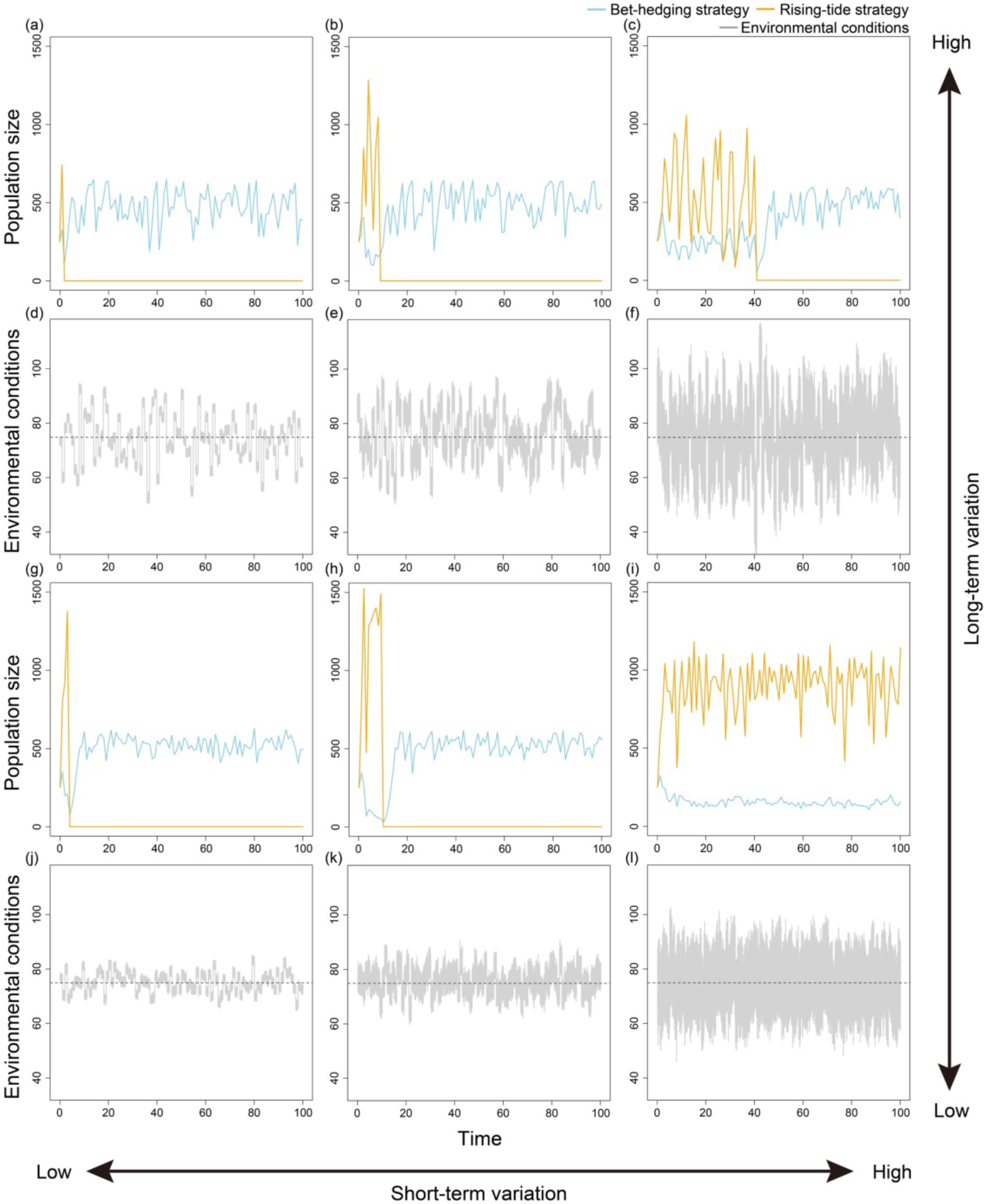
The selection dynamics of the bet-hedging and rising-tide strategies in random continuous environmental conditions with discrete population dynamics. **a-c**, Population dynamics of the bet-hedging and rising-tide strategies with the same parameter spaces described in Fig. 2c-e. **d-f,** Representations of the environmental conditions (i.e. time series) corresponding to panels **a-c. g-i,** Population dynamics of the bet-hedging and rising-tide strategies with the same parameter spaces described in Fig. 2i-k. **j-l,** Representations of the environmental conditions (i.e. time series) corresponding to panels **g-i.** The parameter values of each panel are identical to those in Fig. 2.

The key features of nearly all previously-published non-overlapping generations models (e.g.^1,11,13,23^) are that the fitness consequences of environmental variation within a generation are additive within an organism’s lifetime, but multiplicative among generations. Thus, mathematically, the long-term growth rate of each strategy is determined by its geometric mean of fitness^28-31^. As a result, bet-hedging generalists occur more commonly under a non-overlapping generations scenario because a small absolute fitness in any generation will have a detrimental effect on the long-term growth rate of a strategy. Since our model allows for overlapping generations, birth and death events, and continuous changes in the strength of selection, the rising-tide strategy will be favored by natural selection under a wide range of environmental conditions as long as these specialists are able to encounter their optimal environment (i.e. good years) frequently enough so that they can sustain through adverse environmental conditions (i.e. bad years), even when they achieve lower fitness than under optimal conditions.

In summary, we have identified a novel form of behavioral adaptation to fluctuating environments: the rising-tide strategy. Although both the rising-tide and the bet-hedging strategies are possible when environmentally-driven fitness variation is high, the rising-tide strategy will be favored when mean fitness is higher and the bet-hedging strategy when mean fitness is lower. This difference in which strategy will be favored is determined by the grain of environmental variation, or the frequency of good and bad years within unpredictable environments. Previous theories have provided a heuristic understanding of adaptation to fluctuation environments by considering the mean and variance of fitness associated with the change of environmental conditions in species with non-overlapping generations and populations of fixed size. Our model further shows that under more biologically realistic scenarios involving species with overlapping generations and populations that fluctuate in size as environmental conditions change, even with the same mean and variance in environmental fluctuation, different temporal distributions of environmental conditions can lead to distinct biological adaptations associated with behavioral specialization or generalization. Thus, we provide a general framework for understanding the impacts of different temporal scales of environmental fluctuation on organisms with either overlapping or non-overlapping generational life histories. Future studies must move beyond viewing environmental variation as discrete classes of coarse-(among-generation) versus fine-grain (within-generation) and begin to investigate the existence of a potentially rich suite of adaptations to diverse environmental scenarios—those that vary in intensity, frequency, and duration—in an ever-changing world. Ultimately, our study not only helps bridge the apparent gap between theoretical and empirical studies of biological adaptation in a volatile world, but it also links under a synthesized theoretical framework seemingly distinct fields, such as macrophysiology^32^, species distribution modeling^14,33^, and social evolution^10,18,34^.

## Methods

### Discrete environment model (good and bad years, rising-tide and bet-hedging strategies)

The simplest way to capture this concept is to use two sets of competitive Lotka-Volterra equations (eqns. 1 and 2) with different equilibrium points. We constructed (i) an individual-based model with probabilistic algorithms to determine the upcoming environment and the birth and death events (i.e. a stochastic continuous time model with random environment setting, Fig. 1), and (ii) a deterministic model with traceable environmental change (i.e. deterministic continuous time model with periodical environment setting, Supplementary Fig. 1).

We use a dynamic time step (similar to Gillespie’s algorithm, see Supplementary Fig. 2) in the individual-based model,

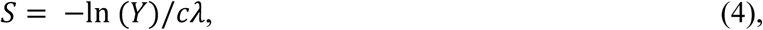

where *S* is the waiting time till next event, *λ* is the sum of all event rates of the differential equations (i.e. birth and death events, Supplementary Fig. 2), *Y* is a random number providing stochasticity to the waiting time, and c is a coefficient for adjusting the timescale. Whenever *S* is determined, a random event occurs according the rate of each events such that a new individual is born, or a living individual dies. As *S* deceases with increasing *λ*, this design flexibly changes the frequency of events based on the current population size.

How the environmental conditions are chosen is another crucial feature of these models. In the individual-based model, after one year has passed, the type of the next year is determined by the probability of good year through a random number draw. On the other hand, we assume good and bad years alternate in the deterministic model but the duration of each changes proportionally to the probability of good year (see supplementary information for more details of the deterministic model).

### Continuous environment model (performance curve, bet-hedging generalist and rising-tide specialist)

To generalize our results and compare them to previous model^15^, we derived the continuous environment model (eqn. 3). Specifically, the carrying capacity of a strategy (*i*) follows a Beta function,

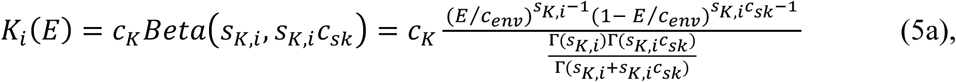

where the shape parameters (*s*) determine the width and height of the performance curve, and the coefficients control the scale (*c_env_*,*_K_*) or skewness (*c_csk_*) of the curve. Further, Γ represents the gamma function, where *Γ*(*n*)=(*n*−1)!. Similarly, birth and death rate follow Beta functions with the same skewness (*c_sk_*) and the same range of environmental conditions (*c_env_*),

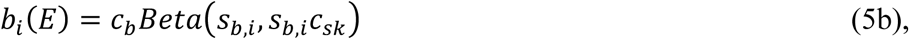

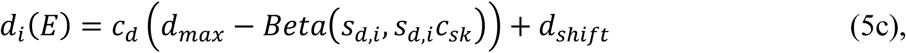

Because the skewness of the three parameters are equal, the mean of each beta function is the same (i.e. *c_env_*(1/1+*c_sk,i_*)). In other words, the only difference in the performance curves among all competitors is the shape parameter, making one strategy more like a bet-hedging generalist or a rising-tide specialist (Fig. 1a). Through this design, the competition in dynamics can respond to the various amounts of change in environmental conditions (e.g. changes in temperature or precipitation).

We have described how parameter values and the population dynamics of each strategy respond to various forms of environmental conditions. Next we determine how environmental conditions fluctuate through time. Since variation may occur in either the long-or short-term, we divide time into phases and episodes, where one phase consists of several episodes. In each time unit, the environmental conditions follow a symmetric Beta distribution,

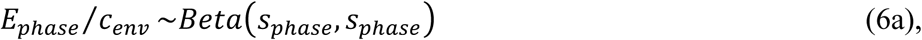

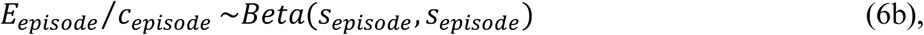

where the scaling coefficients define the range of environmental conditions, and the shape coefficients define the distributions and variation in environmental conditions. Hence, the current environmental conditions are the sum of two sampled values: one from the distribution of long-term variation (controlled by *s_phase_*), and another from the distribution of short-term variation (controlled by *s_episode_*),

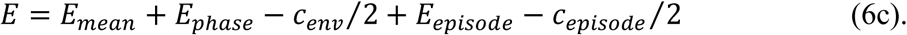

We validate the temporal scale of environmental fluctuations by the fast Fourier transformation (FFT), showing increasing long-term variation enhances the amplitude of low frequency components and increasing short-term variation enhances the amplitude of high frequency components (Supplementary Fig. 3).

Based on the continuous environmental settings, we derive two models using this framework: (i) a continuous; and (ii) a discrete population dynamics model. The continuous population dynamics model (i.e. a stochastic continuous time model with random continuous environmental settings) has the same demographic property as the deterministic environment models, where growth parameters (e.g. *b_s_* and *d_s_*) change instantly with the change of environmental conditions and the population dynamics follows eqns. 3a and 3b. On the other hand, suppose there are *n* episodes in one phase, the discrete population dynamics model (i.e. a stochastic discrete time model with random continuous environment setting) takes arithmetic mean from several parameters by the end of each phase,

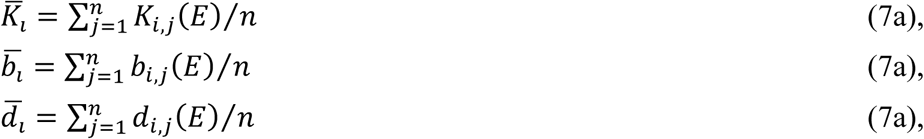

where *i* specifies the strategy and *j* denotes the index of episodes. Hence, the growth parameters are updated once per phase, and the population dynamic follows

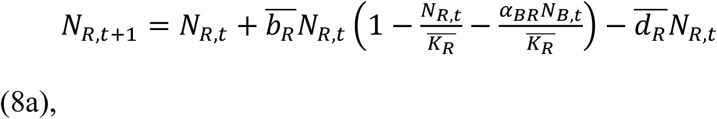

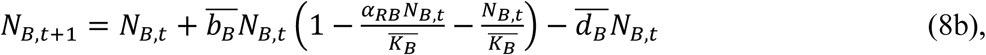

where *t* stands for the index of phases. Because of the process of taking averages, the meaning of short-term variation is no longer the smaller temporal scale of environmental changes. Instead, it expresses a finer type of variation (i.e. finer grain) that potentially deviates from the environment of each episode (Fig. 3).

## Data Availability

All simulated data was generated using the C language. All code used for this study is available at https://github.com/mingpapilio/Codes_RisingTide.

## Author Contribution

S.-F.S. conceived the idea. M.L., W.-C. L. and S.-F.S. constructed the models. M.L., D.D.R. and S.-F.S. designed the study, analyzed the data, and wrote the paper.

## Acknowledgments

We thank Joshua B. Plotkin for the helpful comments for the earlier version of this paper. S.-F.S. was supported by Academia Sinica (Career Development Award and Investigator Award, AS-IA-106-L01) and Minister of Science and Technology of Taiwan (S.-F.S., 1002621-B-001-004-MY3, and 104-2311-B-001-028-MY3). D.R.R. was supported by the US National Science Foundation (IOS-1257530, IOS-1656098).

## Supplementary information

### The deterministic continuous time model with periodical environment setting

Here we reduce the stochasticity of the stochastic Lotka-Volterra model in two ways to make it deterministic and more predictable. First, we calculate the population size from differential equations without using the individual-based model, and then we let good and bad years alternate one after another without randomly picking the type of subsequent year, as in the case in the stochastic continuous time model with random environment setting (i.e. Fig. 1). Hence, a good year (eqns. 1a and 1b) starts after a bad year ends (eqns. 2a and 2b). In addition, we assume that the duration of good and bad years changes proportionally to the probability of a good year occurring, while the sum of one good year and one bad year remains the same.

The simulations of the deterministic model (Supplementary Fig. 1a-c) show qualitatively similar results to the individual-based model (Fig. 1c-e). That is, the rising-tide strategy is more adaptive when good years are more frequent, whereas the bet-hedging strategy is more adaptive when bad years are more frequent. Additionally, phase-plane analysis validates our finding that fitness variation is greater in the rising-tide strategy (Supplementary Fig. 1d-f, amplitude of oscillation on x-axis) relative to the bet-hedging strategy (Supplementary Fig. 1d-f, amplitude of oscillation on y-axis). Thus, these results of the deterministic model are consistent with those of our stochastic model.

**Supplementary Figure 1.**
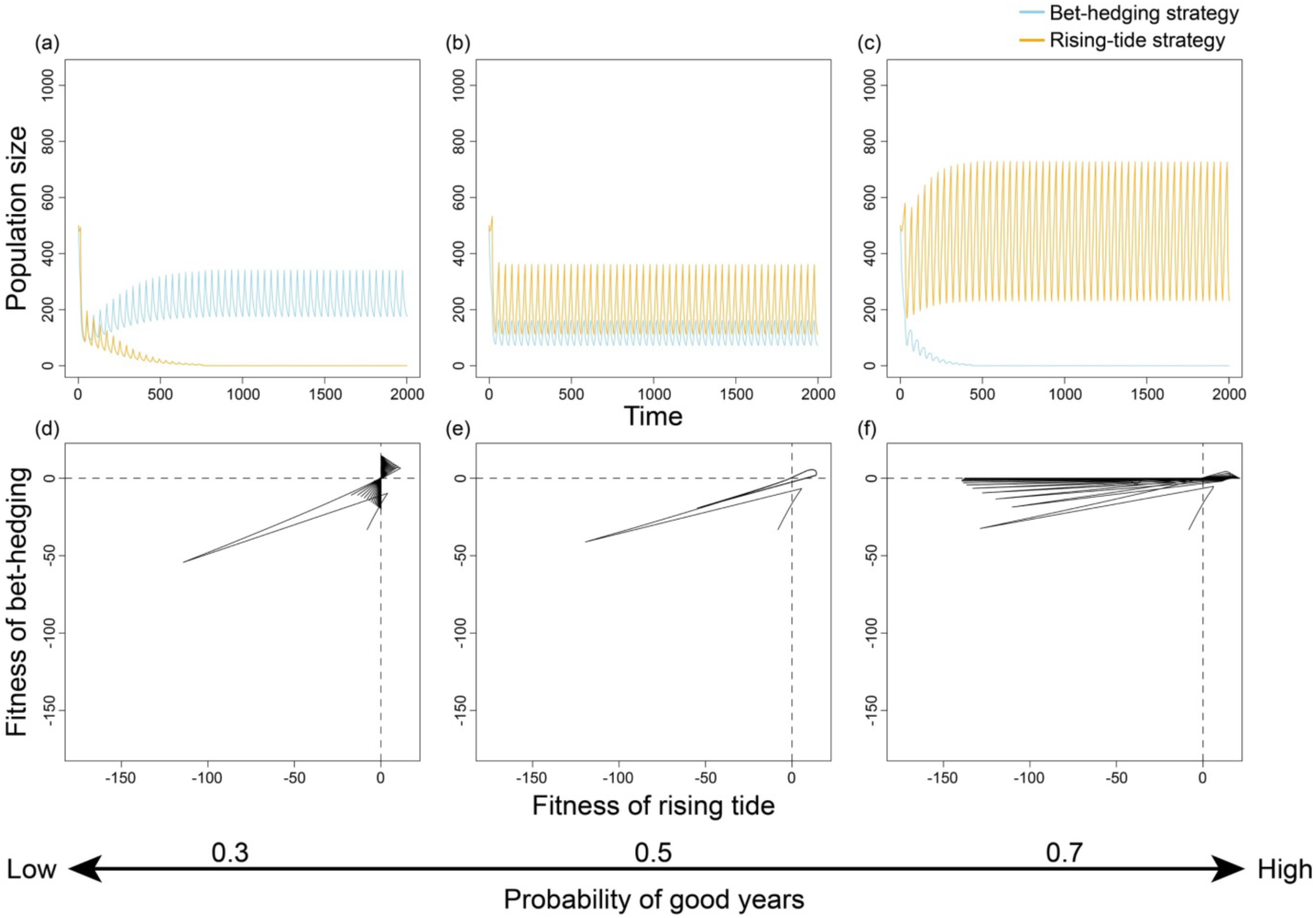
The deterministic continuous time model with periodical environment setting shows qualitatively similar results as the discrete-strategy model (corresponding to Fig. 1). **a-c**, Population dynamics of the bet-hedging (blue lines) and rising-tide strategies (orange lines) in the deterministic model. **d-f**, Phase-plane analysis (i.e. the time series of per capita growth rates) shows the fitness interactions of the bet-hedging and rising-tide strategies.

**Supplementary Figure 2.**
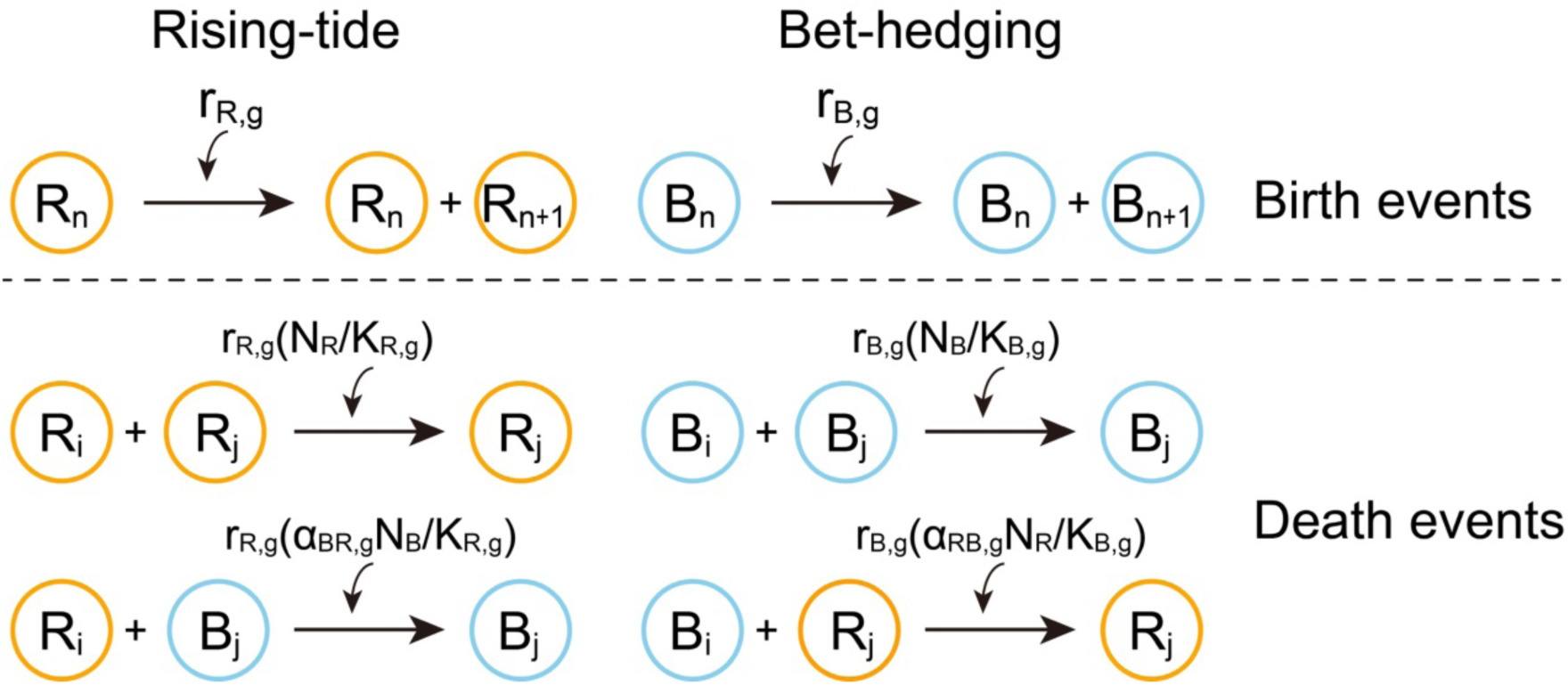
Schematic view of the events in good years for the individual-based model under discrete environmental conditions (corresponding to Fig. 1). Each circle indicates an individual of the rising-tide (orange) or the bet-hedging strategy (blue). Arrows describe the occurrence of birth or death events. The polynomials above the arrows are the probability of each event occurring in the model.

**Supplementary Figure 3.**
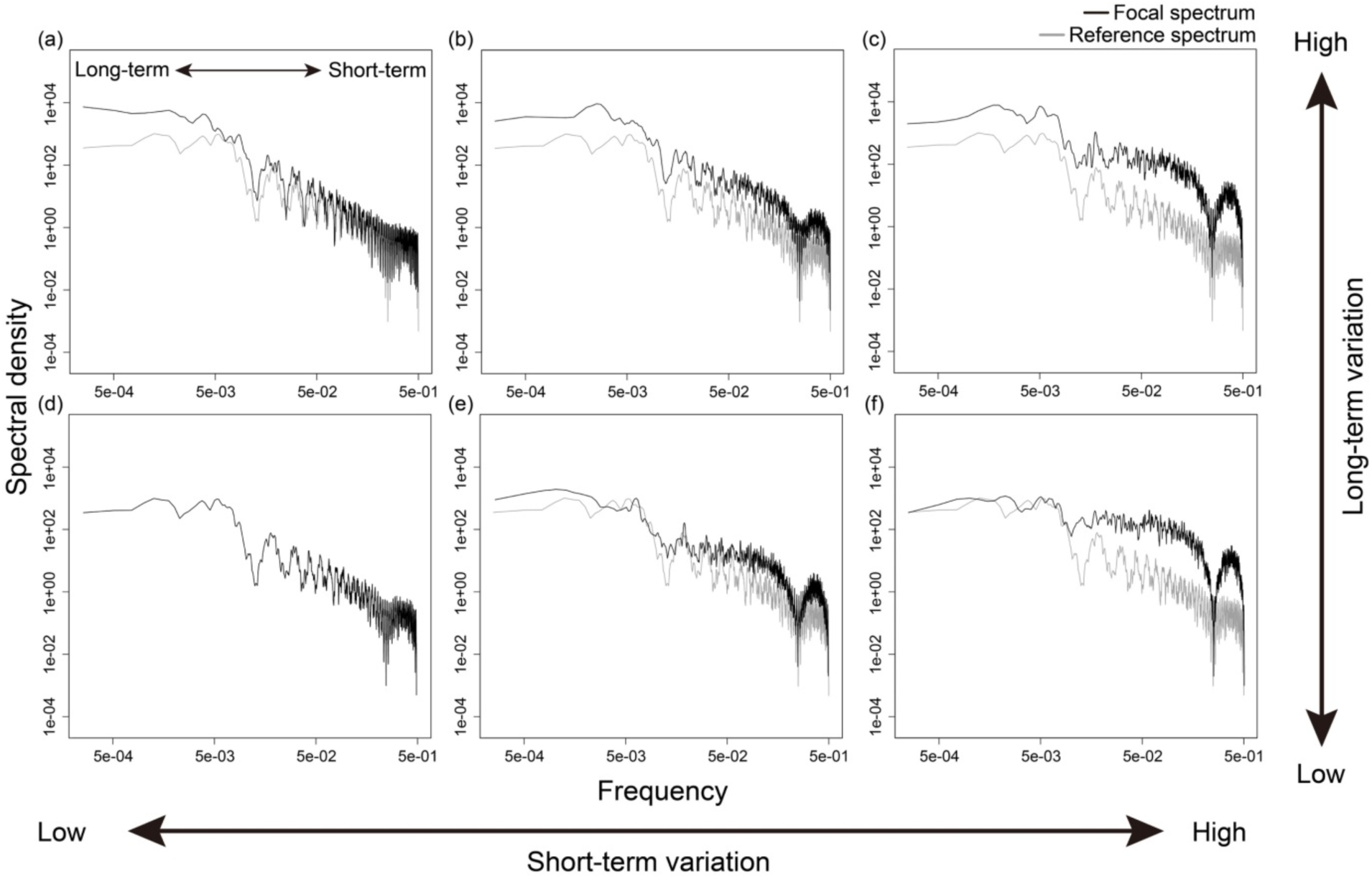
Frequency analysis of environmental conditions in a stochastic continuous time model with continuous environmental settings (corresponding to Fig. 2). **a-f**, We use the fast Fourier transform to understand frequency components in the time series of environmental conditions. In other words, the environmental fluctuations are described by the contributions of each frequency of oscillation (black lines). For comparison, we use the most stable environment (panel *d*) as a reference (gray lines) in all panels. The increasing short-term variation enhances the contribution of high-frequency regions, whereas increasing long-term variation enhances the contribution of low-frequency regions.

**Supplementary Table 1.**
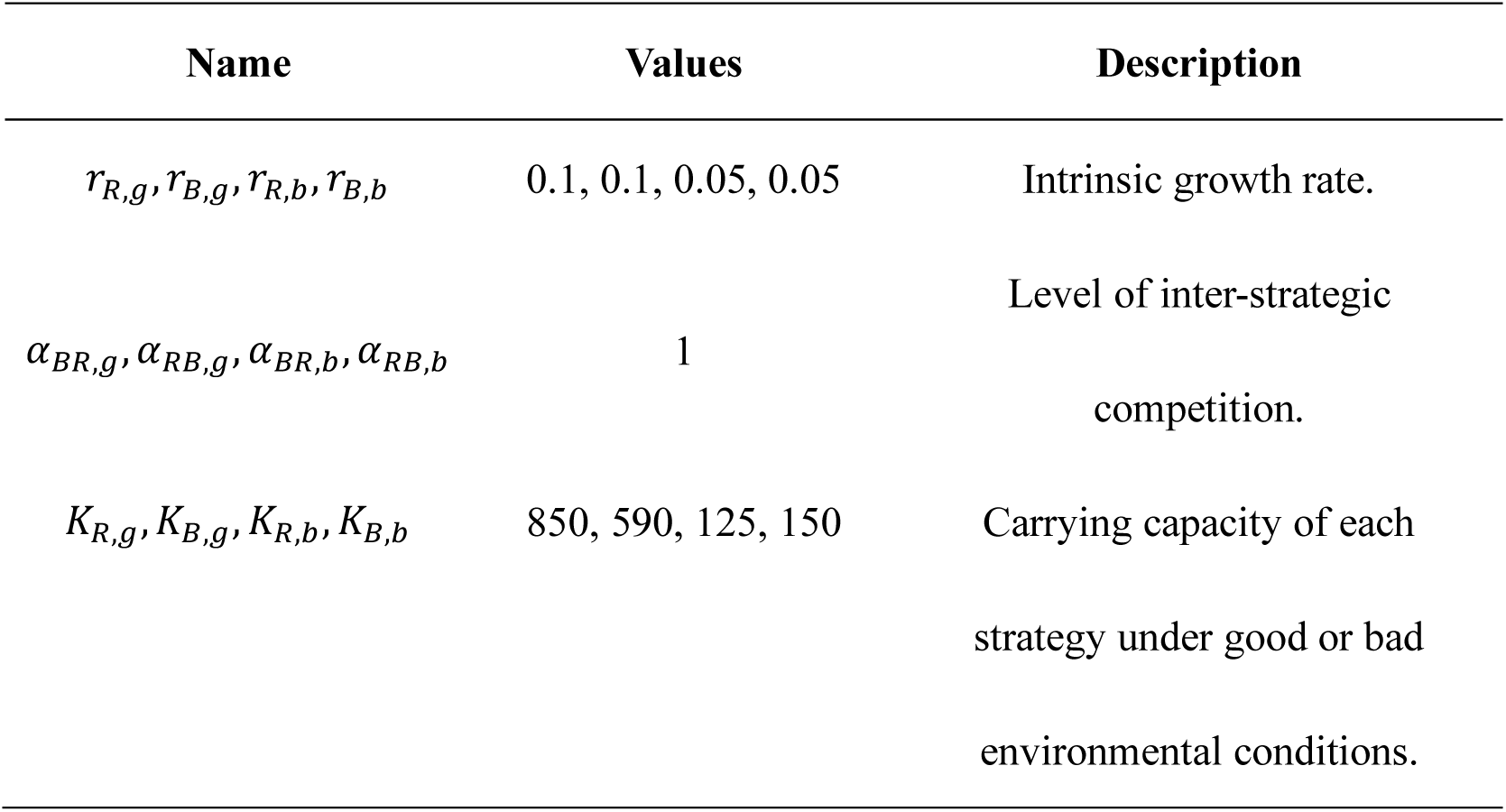
**List of parameters in discrete environment model.** These values are used in the simulations of the individual-based model (Fig. 1) and the deterministic model (Supplementary Fig. 1).

| Name | Values | Description |
| --- | --- | --- |
| $r_{R,g}, r_{B,g}, r_{R,b}, r_{B,b}$ | 0.1, 0.1, 0.05, 0.05 | Intrinsic growth rate. |
| $\alpha_{BR,g}, \alpha_{RB,g}, \alpha_{BR,b}, \alpha_{RB,b}$ | 1 | Level of inter-strategic competition. |
| $K_{R,g}, K_{B,g}, K_{R,b}, K_{B,b}$ | 850, 590, 125, 150 | Carrying capacity of each strategy under good or bad environmental conditions. |

**Supplementary Table 2.** **List of parameters in continuous environment model.** These values are used in the simulations of the continuous (Fig. 2) and the discrete population dynamics model (Fig. 4).

| Name | Values | Description |
| --- | --- | --- |
| $c_K, c_b, c_d$ | 250, 0.5, $10^{-6}$ | Scaling coefficients of population dynamics parameters (carrying capacity, birth rate, and death rate, respectively). |
| $s_{K,i}, s_{b,i}, s_{d,i}$ | 30 (specialist rising-tide), 4.5 (generalist bet-hedging) | Shape coefficient of population dynamics parameters (which determine the widths of performance curves) where larger values indicate narrower distributions. |
| $c_{sk}$ | 1/3 | Skewness of Beta functions, where performance curves follow Beta ($\alpha = s_{j,i}, \beta = c_{sk}s_{j,i}$) and j and i stand for parameters (K, b, d) and strategies (G, S). |
| $\alpha_{GS}, \alpha_{SG}$ | 0.4 | Inter-specific competition coefficient. |
| $c_{episode}, c_{phase}$ | 60, 100 | Scaling coefficients of short- and long-term variation in environmental conditions. |
| $d_{shift}$ | 0.0 | Mortality adjustment for specialist strategy. |
| $s_{episode}, s_{phase}$ | 17*, 10* | Shape coefficient of environmental variation. Episode stands for long-term variation and phase stands for short-term variation. |
| $m_{phase}, m_{generation}$ | 20, 500 | Number of episodes in a phase, and number of events in single generation. |
| $d_{max}$ | $\max [Beta(s_{d,i}, s_{d,i}c_{sk})]$ | Maximum value of death Beta functions among all environmental conditions. |

\*These values are variables.

